# A species-specific lncRNA modulates the reproductive ability of the Asian tiger mosquito

**DOI:** 10.1101/2022.07.04.498273

**Authors:** Alexandros Belavilas-Trovas, Maria-Eleni Gregoriou, Spyros Tastsoglou, Olga Soukia, Antonis Giakountis, Kostas Mathiopoulos

**Affiliations:** Laboratory of Molecular Biology & Genomics, Department of Biochemistry & Biotechnology, University of Thessaly, Larissa, Greece; DIANA-Lab, Department of Computer Science and Biomedical Informatics, University of Thessaly, Lamia, Greece; Hellenic Pasteur Institute, Athens, Greece

**Author notes:** **Correspondence:** Kostas Mathiopoulos. Insect Pest Control Laboratory, Joint FAO/IAEA Centre of Nuclear Techniques in Food and Agriculture, Department of Nuclear Sciences and Applications, IAEA Laboratories, Seibersdorf, Austria. These authors have contributed equally to this work (Maria-Eleni Gregoriou, Spyros Tastsoglou).

**Keywords:** Aedes albopictus, Tiger mosquito, RNAi pest control, lncRNAs, (long non-coding RNAs), species-specific control

## Abstract

Long non-coding RNA (lncRNA) research has emerged as an independent scientific field in recent years. Despite their association with critical cellular and metabolic processes in plenty of organisms, lncRNAs are still a largely unexplored area in mosquito research. We propose that they could serve as exceptional tools for pest management due to unique features they possess. These include low inter-species sequence conservation and high tissue specificity. In the present study, we investigated the role of ovary-specific lncRNAs in the reproductive ability of the Asian tiger mosquito, *Aedes albopictus*. Through the analysis of transcriptomic data, we identified several lncRNAs that were differentially expressed upon blood feeding; we called these genes Norma (<u>NO</u>n-coding <u>R</u>NA in <u>M</u>osquito ov<u>A</u>ries). We observed that silencing some of these Normas resulted in significant impact on mosquito fecundity and fertility. We further focused on Norma3 whose silencing resulted in 43% oviposition reduction and 53% hatching reduction of the laid eggs, compared to anti-GFP controls. Moreover, a significant downregulation of a neighboring (∼100 Kb) mucins cluster was observed in smaller anti-Norma3 ovaries, indicating a potential mechanism of in-*cis* regulation between Norma3 and the mucins. Our work constitutes the first experimental proof-of-evidence connecting lncRNAs with mosquito reproduction and opens a novel path for pest management.

## 1 INTRODUCTION

The remarkable progress of next-generation sequencing and genomics technologies that took place during the past twenty years revealed an unexpected world of transcribed, non-coding (nc) genomic elements that by far exceed in numbers the protein-coding transcripts (Claverie, 2005). Long non-coding RNAs (lncRNAs) represent one class of functional ncRNA transcripts, characterized by species specificity and tissue-specific expression patterns. LncRNA transcripts are longer than 200 nucleotides, they are mainly transcribed by unique genes and most of them are subject to post-transcriptional modifications (splicing, poly-A tail, C-cap), although they have limited or no protein-coding potential. Various research studies in eukaryotic organisms have highlighted the role of lncRNAs in essential biological processes and different modes have been proposed regarding their action. These modes include, but are not limited to, (i) guiding or decoying transcription factors, (ii) acting as scaffolds for chromatin modifying complexes, (iii) functioning as sponges for miRNAs, (iv) regulating post-transcriptional mRNA modifications [reviewed in (Marchese et al., 2017)].

Due to the absence of coding capacity, lncRNAs demonstrate a notable lack of nucleotide sequence conservation even among closely related species, which results in a high number of unique, species or genus specific lncRNAs across eukaryotic organisms (Pang et al., 2006; Bhartiya and Scaria, 2016). This low sequence conservation along with the aforementioned multifunctionality provide extraordinary difficulties in the development of computational tools that would predict lncRNA targets or potential modes of action. Nonetheless, despite this sequence variation, lncRNAs from different organisms may exhibit structural or functional conservation, highlighting their conserved role in essential biological pathways (Ponjavic et al., 2007; Diederichs, 2014; Tavares et al., 2019). Indeed, unlike other non-coding RNAs (e.g., miRNAs) that hybridize to their targets through sequence complementarity patterns, the functionality of lncRNAs mainly results from their secondary structure motifs (Kino et al., 2010; Ding et al., 2014; Chillón and Pyle, 2016; Smola et al., 2016).

The high sequence divergence of lncRNAs renders them as ideal candidates for the development of species-specific population control approaches against organisms of interest. Achieving species-level specificity is a point of major importance for the development of novel pest management approaches especially against insect pests, such as mosquitoes. Current insect control approaches are mainly based on chemical insecticides that pose serious threats to public health and biodiversity due to their neurotoxic action against other species, either mammals (including human) (Costa et al., 2008) or off-target insects (Desneux et al., 2007). Beneficial insects, contributing vitally to the stability of ecosystems and to agriculture, are severely harmed by the main classes of pesticides, even by those that are considered safe for humans. This issue arises due to the lack of species-specific mechanisms underlying insecticide action. Both organophosphates and carbamates target acetylcholinesterase (AChE) which is conserved among insects (Kwong, 2002), while pyrethroids target conserved voltage-sensitive sodium channels (Soderlund, 2010). Moreover, neonicotinoid pesticides which are considered non-toxic for vertebrates, act as agonists of nicotinic acetylcholine receptors (nAChRs). However, all insects are vulnerable to these pesticides due to the conserved nAChRs sequences and their critical role in signal transduction (Simon-Delso et al., 2015). In addition to the lack of species-specificity in the currently used chemical insecticides, there is also the huge and long-standing problem of insecticide resistance. The way out of this unfavorable situation is the development of novel pesticides that target alternative gene classes that could lead to more effective pest management (Sparks et al., 2021).

The Asian tiger mosquito, *Aedes albopictus*, is also the target of various control approaches, as it is a cosmopolitan vector of several lethal arboviruses, including dengue, Zika and chikungunya (Martinet et al., 2019). *Ae. albopictus* emerged from the tropical and sub-tropical regions of south-east Asia and rapidly expanded throughout the world due to its exceptional ability to adapt in different environmental conditions (Benedict et al., 2007). Its vectorial status arose especially after its connection with the major outbreaks of Chikungunya virus in 2005-07 in La Reunion (Pialoux et al., 2007) and in 2007 in Italy (Rezza et al., 2007; Angelini et al., 2008), while it was also associated with the first autochthonous cases of dengue fever in France in 2010, 2013 & 2015 (La Ruche et al., 2010; Marchand et al., 2013; Succo et al., 2016). It is certain that in the coming years its expansion will continue, and estimates indicate that by 2050 half of the world population will be exposed to disease-spreading mosquitoes, such as *Ae. albopictus*, due to climate change and global warming (Kraemer et al., 2019).

In the present study we explore the potential of targeting lncRNAs to control insect populations. LncRNAs could be used as species-specific molecular targets for the development of next-generation pesticides (e.g., RNAi pesticides (Fletcher et al., 2020)) or be part of the rapidly growing synthetic biology systems, such as SIT and gene drives (Caragata et al., 2020). The present study focuses on the investigation of lncRNAs that are related to *Ae. albopictus* reproduction, due to the significance of reproduction in population suppression approaches. The reproductive process in females is triggered by the consumption of a blood meal (BM) which activates a cascade of metabolic signaling pathways that lead to the development and production of eggs. We sought to identify lncRNAs that influence the reproductive process, aiming at the disruption of oogenesis and the reduction of mosquito fecundity and fertility.

## 2 MATERIALS AND METHODS

### 2.1 Mosquito rearing

An *Aedes albopictus* laboratory line was established from wild mosquitoes which were collected from the region of Thessaly, as described elsewhere (Ioannou et al., 2021), and were reared in the insectary facility of the Department of Biochemistry & Biotechnology at the University of Thessaly. Adult mosquitoes were reared at 26±1°C with a relative humidity of 60-70%, under a 14h:10h light/dark photoperiod. They were fed with 10% sucrose solution, while female mosquitoes were blood-fed (BF) from a human arm to initiate their gonotrophic cycles.

### 2.2 RNA extraction, reverse transcription and Real-Time PCR

Ovaries and other tissues were dissected under the microscope and their total RNA was extracted using Extrazol (BLIRT S.A., Gdańsk, PL), according to the manufacturer’s instructions. The integrity of the RNA was assessed by agarose gel electrophoresis. Total RNA was treated with DNase I (Thermo Fisher Scientific, Waltham, MA, USA) and 1μg of RNA was reverse transcribed to cDNA by using oligo-dT primers and MMLV-RT (Invitrogen, Waltham, MA, USA), according to the manufacturer’s instructions. Each biological replicate corresponds to tissues collected from individual mosquitoes. We preferred to study tissues from individual mosquitoes, rather than pooling them together, in order to be able to assess within population variability. All qPCR assays were performed in CFX96 Real-Time Thermal Cycler (Bio-Rad, Hercules, CA, USA). All amplifications were performed with two technical replicates and the relative gene expression was analyzed by using the 2^-ΔΔCt^ method (Livak) through the CFX Manager™ software. Specific primers to amplify the genes identified by the transcriptomic analysis were designed using PrimerQuest™Tool. Their target specificity was verified through Primer-BLAST (Ye et al., 2012) against the Refseq mRNA database of *Aedes albopictus*. Primers that lacked any sequence homology to other transcripts were selected. Two endogenous ribosomal housekeeping genes (RpL32, RpS17) were used for the normalization of the results (Dzaki and Azzam, 2018). The average expression stability value (M-value) of the reference genes for each biological sample was determined and samples that exhibited M-value <0.5 and Coefficient of Variance <0.25 were accepted for further analysis. Primers for qPCR are shown in Supplementary Table S2.

### 2.3 In Vitro Double-Stranded (ds) RNA Synthesis-dsRNA treatment

Target templates for *in vitro* transcription were generated using gene specific primers with the respective recognition site for T7 RNA polymerase designed by the eRNAi web platform (Horn and Boutros, 2010). Thermodynamic parameters of the primers were tested through the web platform OligoAnalyzer (https://eu.idtdna.com/calc/analyzer). We selected primer sets that exhibited a minimum ΔG value of -9kcal/mole for homodimer and heterodimer formation, while their hairpin structures did not exceed the primer Tm-10°C. Their target specificity was assessed by Primer-BLAST (Ye et al., 2012) against the Refseq mRNA database of *Aedes albopictus*. dsRNA was synthesized by using the MEGAscript RNAi kit (Ambion, Austin, TX) from a dsDNA template by incubating at 37°C for 16h. dsDNA was produced by PCR with cDNA template and gene-specific primers with the T7 RNA polymerase promoter (TAATACGACTCACTATAGGG) attached to their 5’ end. dsRNA was purified with standard phenol/chloroform protocol, its integrity was measured by agarose gel electrophoresis and its concentration was quantified by the Q3000 Spectrophotometer (Quawell, San Jose, CA). Green fluorescent protein (GFP) dsRNA was used as a control. For silencing with each one of the Norma genes, inseminated *Ae. albopictus* adult females were used. The mosquitoes were collected five days after eclosion and were anesthetized on CO_2_. Then, 64.4nl of dsRNA solution (5,000ng/μL) was injected into their thorax using the Nanojet II microinjector (Drummond, Birmingham, AL), under a Leica™ stereoscope. At least 25 individual females (i.e., biological replicates) were used for silencing with each Norma gene. Two controls were used for comparison: one was non-injected females (untreated) and the other was GFP-dsRNA injected females (anti-GFP). Both control populations were reared under the same conditions with the Norma-dsRNA injected ones. Primers for dsRNA production are shown in Supplementary Table S2.

### 2.4 Phenotypic assays

Five-day-old mosquitoes injected with either GFP-dsRNA or Norma3-dsRNA were transferred to Bugdorm™ 17.5×17.5×17.5cm cages (MegaView Science Co., Talchung, Taiwan) immediately after injection, where they were fed with 10% sucrose solution for 24h to recover. An additional sample of same age, non-injected mosquitoes (untreated) was also maintained. The mosquitoes were left to starve for 12h and subsequently they were blood-fed (36h post-injection). All blood meals took place in the same timeframe, between morning and noon, to avoid the influence of circadian variability among the replicates. Fully engorged, blood-fed mosquitoes were reared in the presence of 10% sucrose.

#### 2.4.1 Ovarian measurement

Ovaries were dissected 60h post blood meal by detaching the soft cuticle between the fifth and sixth abdominal segment and pulling the terminal segments with fine forceps onto a drop of phosphate buffered saline (PBS). Pictures of the ovaries were captured with a Leica™ stereoscope (Leica Microsystems, Wetzlar, Germany) and the length of the long axis of the ovarian follicle was measured by FIJI software (Schindelin et al., 2012).

#### 2.4.2 Oviposition

Four days after blood feeding individual mosquitoes were placed in polystyrene fly vials that contained a filter paper attached to a wet cotton ball and were left into the vial for two days in the insectary to lay their eggs. Additional moisture was added regularly to the vials to keep the filter paper wet. Two days later, mosquitoes were removed from the vials and the total number of eggs deposited by each individual mosquito was counted under a stereoscope. Mosquitoes that deceased during the process were excluded from the analysis.

#### 2.4.3 Hatching assay

Laid eggs were dried and then stored into sealed petri dishes that contained a wet cotton ball as a source of humidity. Each petri dish hosted the eggs laid by one individual mosquito. Three days (72h) post egg laying, 30ml of hatching broth, that was prepared as described elsewhere (Maïga, 2017), were added into each petri dish. Eggs that were stored inside the petri dishes were incubated with the hatching broth for 14 days. Hatching broth was freshly replaced every five days. The long 14-day period was preferred because shorter periods led to reduced hatching of the eggs, probably due to diapause effect. Emerged larvae from each egg batch were counted daily and were removed from petri dishes. Finally, the hatch rate was estimated as the total number of emerged larvae divided by the total number of laid eggs:

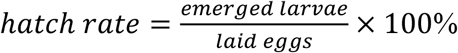

#### 2.4.4 Bleaching assay

The filter papers that contained the eggs that were laid by individual dsRNA-injected females were collected, placed separately into petri dishes and left under moisture for 72h to complete their embryogenesis. Then they were dechorionated in order to visualize their internal segments, based on a clarification methodology described elsewhere (Trpiš, 1970). A fresh Trpiš solution (3gr NaClO_2_, 2ml glacial acetic acid, in 1L distilled H_2_O) was prepared for each experiment and was added into the petri dishes that hosted eggs that were collected from mosquitoes injected with dsRNA against Norma3 or GFP. The eggs were incubated with Trpiš solution for 40 minutes at room temperature and then were immediately washed with PBS and placed gently with a soft brush into a drop of PBS under a Leica™ stereoscope (Leica Microsystems, Wetzlar, Germany) for visualization.

### 2.5 RNA-Seq data pre-processing and primary analysis

Raw FASTQ files from a detailed, publicly available, developmental transcriptomic dataset by Gamez and colleagues were retrieved from the Sequence Read Archive (SRA) study entry SRP219966 (Gamez et al., 2020), using SRA toolkit. Quality assessment, identification of adapters and overrepresented contaminants was performed by employing FASTQC (www.bioinformatics.babraham.ac.uk/projects/fastqc/) and the EMBL-EBI’s Minion application (http://www.dev.ebi.ac.uk/enright-dev/kraken/reaper/src/reaper-latest/doc/minion.html). Quality and adapter trimming of reads was performed with cutadapt. Pre-processed FASTQ files were mapped to the GCF_001876365.2_canu_80X_arrow2.2 *Ae. albopictus* genome assembly utilizing STAR splice-aware aligner (v2.7.9a) (Dobin et al., 2013) using the ENCODE standard options provided in the STAR manual. The respective transcript annotation file, derived from the NCBI Eukaryotic Genome Annotation Pipeline (Gnomon gene predictions), was also provided (--sjdbGTFfile argument) to effectively account for known splice junctions, while the genome index was accordingly parameterized to the read length (--sjdbOverhang 49). Transcript-level expression was calculated with RSEM v1.3.1. (Li and Dewey, 2011).

### 2.6 LncRNA annotation

Accession numbers with the “XR_” prefix, annotated as non-coding RNA (ncRNA) were retained from the GCF_001876365.2 gene models. In order to obtain estimates of the transcripts’ coding potential, CPC2 (Kang et al., 2017) and FEELnc (Wucher et al., 2017) tools were employed. Both tools were executed at default settings (in the absence of any known lncRNAs, FEELnc “--mode=shuffle” method was chosen and the “XM_” (mRNA) transcripts were also provided as input for training). Transcripts were queried locally with BLASTn v2.13.0 (Altschul et al., 1990), at default settings, against Reference RNA Sequences (refseq_rna) from all Hexapoda species (Taxonomy ID: 6960). Each transcript was annotated with respect to the number of species it presented hits to, besides *Ae. albopictus*. FEELnc classifier was also used to characterize all “XR_” transcripts as genic and intergenic, according to their genomic localization and strand, respective to other transcripts. Lastly, RSEM-derived counts from all analyzed datasets in Section 2.6 were TMM-normalized (Robinson and Oshlack, 2010), log_2_-transformed and utilized to obtain *tau* (*τ*) tissue specificity indices (Yanai et al., 2005) (https://github.com/roonysgalbi/tispec). For a transcript *x* with expression *x*_*i*_ across *n* tissues/contexts, *tau* is obtained using its expression normalized by its maximal expression as follows:

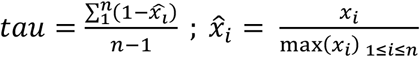

After deriving the global *tau* for a transcript, its per-context specificity fraction was calculated by multiplying *tau* with the maximal-normalized expression metric 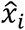. “XR_” transcripts were annotated with their context (i.e., developmental stage, tissue and time-point) *tau* specificity index, as well as per-context fraction. Heatmaps were created using R package pheatmap, using Euclidean distance as a metric and complete-linkage hierarchical clustering where applicable. Annotation results are provided unfiltered in Supplementary Data S1.

### 2.7 Norma3 expression correlation and clustering analysis

For correlation analysis, samples where Norma3 presented non-zero expression (Transcripts-Per-Million, TPM units) were selected and transcripts presenting zero TPM across all samples were filtered out. Pearson correlation testing and False Discovery Rate adjustment of p-values were conducted in R. Clustering analysis was performed by providing TPM values to DPGP (McDowell et al., 2018), which applies Dirichlet Process to nonparametrically determine the optimal number of expression trajectory clusters and Gaussian Process to model the trajectories of expression through time. Following DPGP recommendations, a limited list of transcripts was subjected to clustering analysis instead of the whole transcriptome. For that purpose, read counts from post blood-meal (PBM) ovary samples (12h, 24h, 36h, 48h, 60h and 72h) were imported in R. In the absence of replicates, 12-24h, 36-48h and 60-72h time points were grouped into “early”, “middle” and “late” groups respectively. EBSeq (v1.30.0) (Leng et al., 2013) was utilized to assign transcripts to patterns of differential expression, at a relaxed posterior probability ≥75%. DPGP was executed at default settings.

### 2.8 Statistical analysis

Data were presented as mean ± SEM of independent biological replicates, unless otherwise noted. Distribution patterns of the samples were evaluated through Shapiro-Wilk test (Shapiro and Wilk, 1965) and those populations that followed normal distribution were analyzed by two-tailed unpaired Student’s t-test. Samples that did not pass normality test were analyzed by nonparametric two-tailed Mann-Whitney U test (Mann and Whitney, 1947). P-values of ≤0.05 were considered as significantly different. All analyses were performed through GraphPad Prism 8 software and all values are displayed in Supplementary Data S3.

## 3 RESULTS

### 3.1 Annotation of predicted non-coding RNA in *Ae. albopictus*

In order to query for *Ae. albopictus* lncRNAs that are potentially implicated in mosquito reproduction, we initiated our computational analysis on the respective NCBI gene models, which contain 8571 transcripts that are predicted to belong to the ncRNA class (“gbkey=ncRNA”). Bioinformatics approaches were adopted to annotate further these RNAs regarding their coding potential, species specificity, genomic localization, as well as their context-expression specificity within *Ae. albopictus*. Coding potential estimates of the predicted ncRNAs were calculated with CPC2 and FEELnc. CPC2 (Kang et al., 2017) is a well-known species-agnostic Support Vector Machine model that makes use of Fickett score, open reading frame (ORF) length and integrity and isoelectric point to predict coding potential. In contrast, FEELnc (Wucher et al., 2017) accepts as input known coding and non-coding transcripts from a species of interest and trains a species-aware coding potential model (Random Forests) using as predictors the ORF coverage of transcripts, k-mer composition (1-, 2-, 3-, 6-, 9-, 12-mers) and transcript length. When known non-coding RNA sequences are not available, under shuffle mode, FEELnc shuffles the provided protein coding sequences but preserves 7-mer frequencies, to create a faux set of non-coding transcripts to use in model training-testing, a method shown to fare adequately in benchmarks against real lncRNAs (Wucher et al., 2017). The FEELnc model output is a score between 0 (i.e., no coding potential) and 1 (coding potential), while its optimal cutoff point that maximizes sensitivity and specificity is calculated by means of 10-fold cross validation. CPC2 annotated 8289 “ncRNA” transcripts as non-coding and 282 as potentially coding. FEELnc classification yielded 8096 and 475 transcripts labeled as non-coding and coding respectively, at an optimal cutoff point of 0.43 (Figure 1A). CPC2 and FEELnc predictions were juxtaposed with each other, highlighting 7900 and 86 transcripts (92% and 1% of total) concordantly classified as non-coding and coding RNA respectively, and identifying 589 predictions (7% of total) that were discrepant between the two approaches (Figure 1B, Supplementary Data S1).

**Figure 1:**
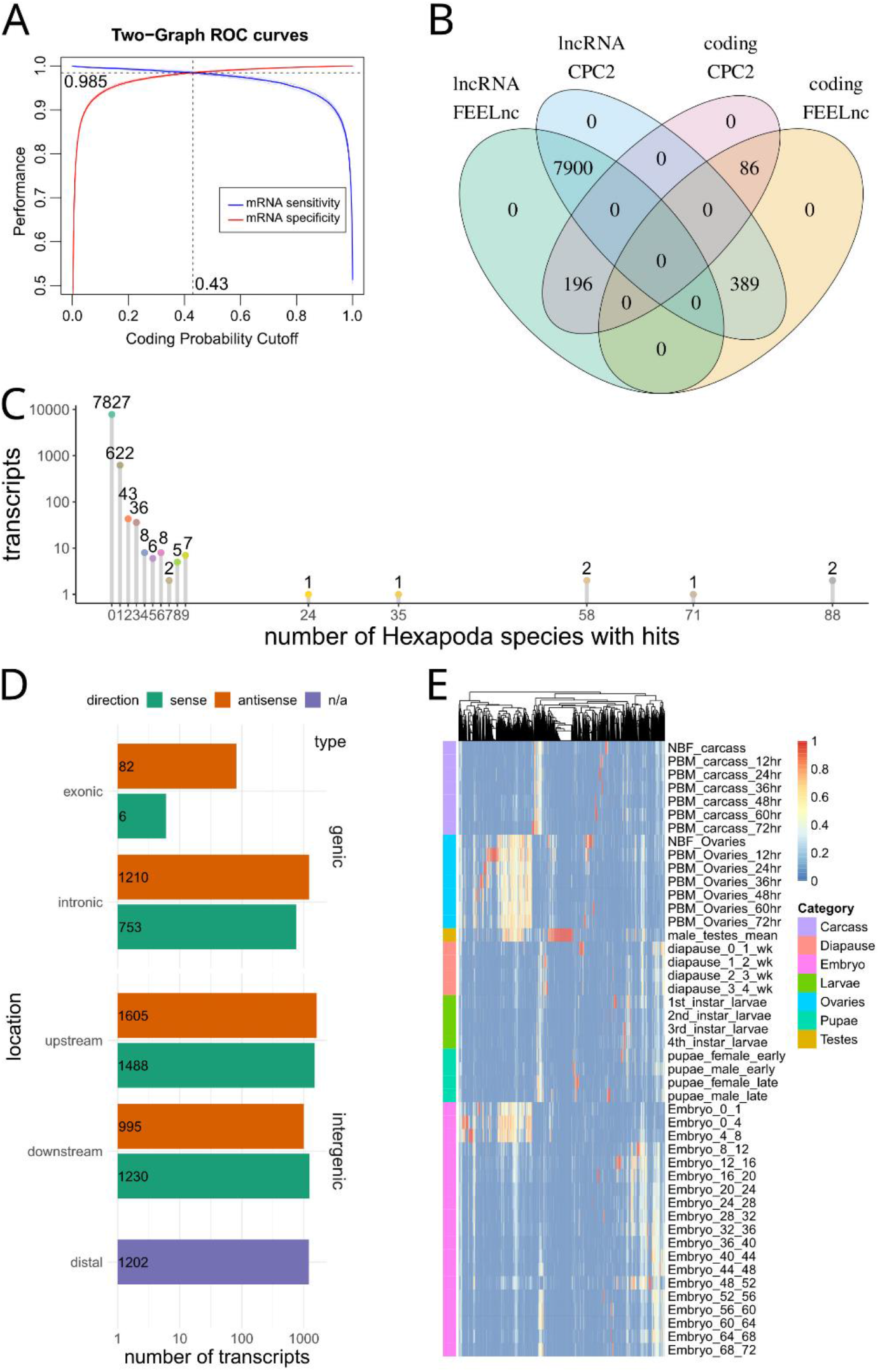
Computational analysis of RNAs predicted to be non-coding via automated NCBI analysis. **(A)** A coding potential prediction model (FEELnc Random Forests) was trained specifically on *Ae. albopictus* sequences. The optimal cutoff to discriminate coding from non-coding sequences was set at the point where sensitivity and specificity was maximized (10-fold cross validation). **(B)** Overlap of coding potential predictions between FEELnc and CPC2 models. **(C)** Depiction of number of transcripts presenting hits to other Hexapoda species. Transcripts are tallied by the number of species in which they presented BLASTn hits (x-axis). **(D)** Genomic localization of lncRNAs, per type (genic/intergenic), subcategory (exonic/intronic/upstream/downstream/distal (i.e., >100kb afar from other transcripts) and strand relative to close/overlapping elements (sense/antisense). **(E)** Heatmap of per-context fractions for all lncRNAs presenting *tau* >0.5.

In order to identify species-specific lncRNAs, transcript homology queries were performed (local BLASTn) against all Hexapoda Reference RNA sequences. Each *Ae. albopictus* “ncRNA” transcript was annotated with respect to the number of other species it presented hits against. Out of the total transcripts, 7827 (91%) were found to exhibit no similarity with RNAs of any other Hexapoda species (Figure 1C).

FEELnc was also utilized to annotate transcripts’ localization with regard to transcripts annotated as protein coding (Figure 1D). In total, 2051 instances (24%) were found to overlap protein coding transcripts in sense (9%) or antisense (15%) orientation. The remaining 76% was divided among intergenic transcripts that presented non-overlapping neighboring genes within 100kb of their loci (5318 transcripts, 62%) and those that were found to exist in genomic regions not harboring other genes (annotated as “distal” intergenic, 1202 instances, 14%).

Finally, in order to obtain expression metrics with regard to these ncRNAs in discrete developmental stages and tissues, a publicly available developmental transcriptomic dataset of *Ae. albopictus* (Gamez et al., 2020) was analyzed from scratch (SRA accession number SRP219966). Briefly, the dataset captured expression estimates from adult ovaries of non-blood-fed (NBF, fed with a 10%-sucrose solution) and post-blood-meal (PBM, at 12-24-36-48-60-72 time-points) insects, carcasses (i.e., female bodies without the ovaries, also NBF and PBM at the same time-points), adult testes, diapause, larvae, pupae and embryo samples at numerous time-points. In the absence of replicates (with the exception of testes which were in duplicate), no attempt to assess differential expression was performed. Instead, log_2_-transformed TMM-normalized counts were used to obtain *tau* indices (Yanai et al., 2005), which have been recently shown to perform consistently as a tissue specificity metric (Kryuchkova-Mostacci and Robinson-Rechavi, 2017). Briefly, *tau* constitutes a per-transcript tissue specificity unit ranging from 0 (no specificity) to 1 (highest specificity), while the fractions of each tissue that contributed to calculation of *tau* can also be obtained. Within the current dataset that is composed of a number of distinct tissues, developmental stages and time-points, we denoted that *tau* be regarded as a context-specificity index. The fractions of 3119 transcripts exhibiting *tau* > 0.5 are presented as a heatmap (Figure 1E) for all available contexts, outlining the existence of numerous instances of intermediate-to-high specificity in ovary contexts (i.e., yellow-to-red transcripts within ovary samples).

We focused our attention on 984 transcripts that were (a) annotated as having no/limited coding potential by both CPC2 and FEELnc, (b) presented no sequence similarity to known transcripts of other Hexapoda and (c) exhibited *tau* >0.5 and *tau fraction* >0.5 in at least one PBM ovary time-point. Aiming to experimentally scrutinize a limited number of lncRNAs, out of this subset we selected 10 ovary-specific lncRNAs that were upregulated upon blood-feeding and presented limited or no expression in other developmental samples (Supplementary Table S1; Figure 2A). We termed these lncRNAs Norma (<u>NO</u>n-coding <u>R</u>NAs in <u>M</u>osquito ov<u>A</u>ries).

**Figure 2:**
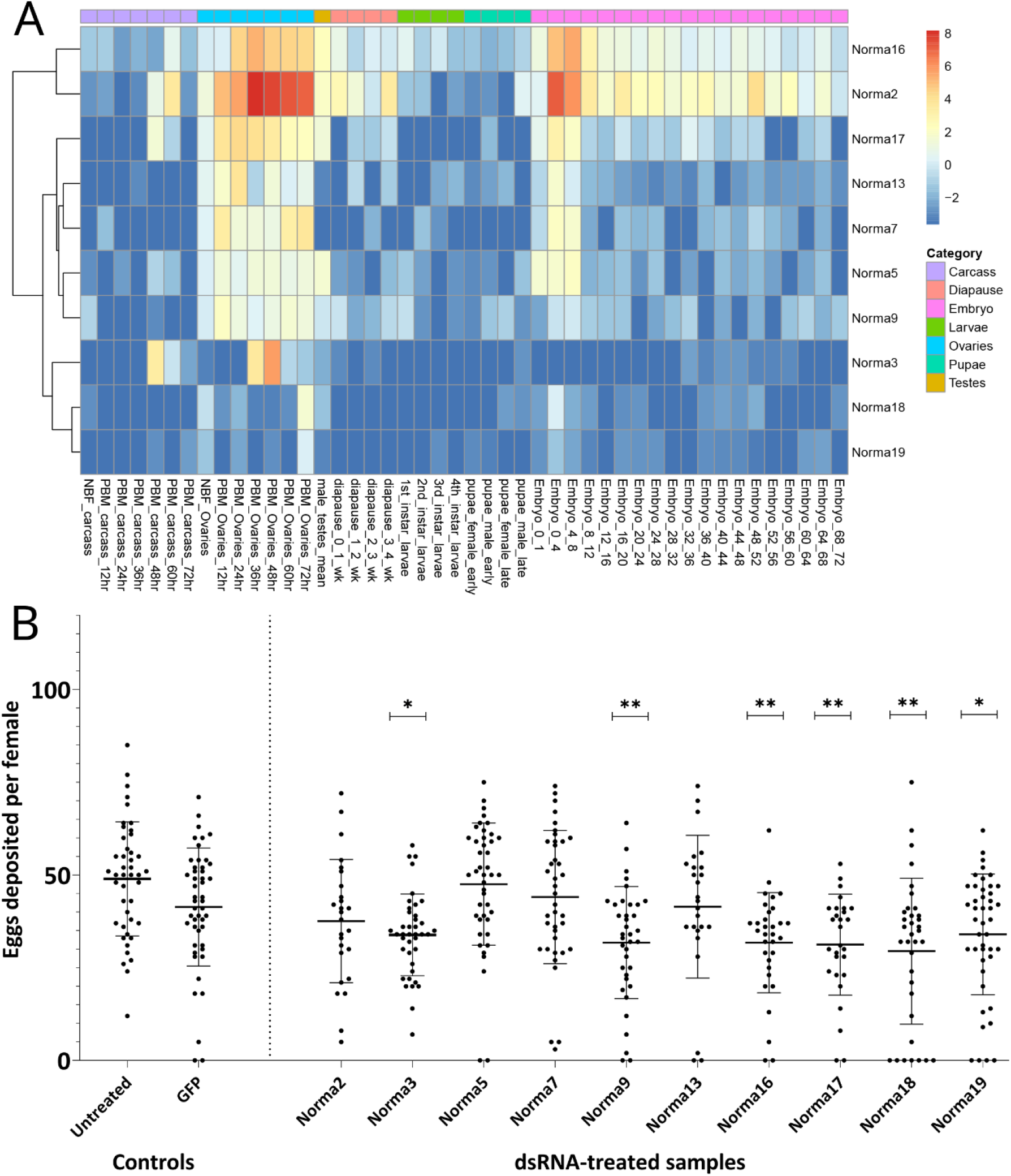
Expression of Norma transcripts across mosquito developmental contexts and oviposition rate change after dsRNA-treatment against them. **(A**) Heatmap of Norma expression levels in carcass, ovary, testis, diapause, larva, pupa and embryo samples from a publicly available transcriptomic dataset by Gamez *et al*.. Expression is depicted as log_2_-transformed TMM-normalized counts. **(B)** Impact of dsRNA-treatment, against each Norma separately, in oviposition rate. Each dot corresponds to the number of eggs that were laid individually by each female mosquito treated with dsRNA against 10 Norma or GFP dsRNA-treated mosquitoes and untreated control. Each sample contained at least 25 mosquitoes (biological replicates) and an unpaired two-tailed student’s t-test was conducted to assess the statistical significance of the results, by comparing the anti-GFP sample (control) with each anti-Norma sample. Treatment with dsRNA against Norma3, 9, 16, 17,18,19 displayed statistically significant differences that may represent the influence of those genes on oviposition. Results are presented as mean ± SD. *:P-value≤0.05, **:P-value ≤0.01

### 3.2 Phenotypic impact of Norma genes silencing

In order to clarify the potential role of the 10 Norma genes, we evaluated their phenotypic impact in reproduction through RNAi silencing. Given their expression pattern, we reasoned that knocking down Norma genes would mostly impact mosquito oviposition. To assess this, we generated *in vitro* dsRNA against each of the ten Norma lncRNAs which we injected into 5-day old inseminated adult females. We then provided a blood meal to the injected mosquitoes and monitored them for oviposition. We counted the number of eggs that were laid by each individual mosquito treated with any of the ten Norma-dsRNA (anti-Norma) and compared oviposition with a GFP-dsRNA sample (anti-GFP). A non-injected, blood-fed sample (untreated) was also present in the assay to monitor the effect of the environmental conditions. We observed that females injected with six different Norma-dsRNAs laid fewer eggs compared to GFP-dsRNA injected control, indicating the potential influence of silencing the corresponding lncRNAs to reproduction (Figure 2B). Specifically, dsRNA against Norma3, Norma9, Norma16, Norma17, Norma18 & Norma19 exhibited statistically significant reduction of oviposition rates compared to the GFP control (P-value≤0.05). We focused our downstream analyses on Norma3 because it also presented some further striking features: an ovary-specific expression profile along with a sharp upregulation pattern in the post-vitellogenic time-points. Other lncRNAs that influenced oviposition (Norma9, Norma16, Norma17, Norma18, Norma19) will be the subject of a future investigation.

### 3.3 Norma3 expression analysis and RNAi-knockdown

Initially, we determined the detailed expression profile of Norma3 by examining its stage- and tissue-related patterns in samples collected from our laboratory mosquito strain. We collected ovaries at the same time-points as the ones that were described in the RNA seq dataset that we analyzed. Specifically, we collected ovaries from NBF and PBM (12h, 24h, 36h, 48h, 60h, 72h) time-points. In addition, we dissected and stored other tissues from the same time-points (carcass, midgut, Malpighian tubes, head). Norma3 exhibited basal expression in NBF and early PBM samples, while its expression abruptly increased >2000-fold in 60h-PBM ovaries, compared to the expression in NBF ovaries, and then significantly dropped to 200-fold in 72h-PBM ovaries, compared to NBF (Figure 3A). At the same time-point (60h PBM), Norma3 displayed basal expression in all other examined tissues (carcass, midgut, Malpighian tubes, head) (Figure 3A). The expression pattern was in accordance with publicly available RNA seq data (Gamez et al., 2020). Subsequently, we estimated the efficiency of Norma3 silencing in the ovaries of female mosquitoes injected with Norma3-dsRNA (anti-Norma3), compared with mosquitoes injected with GFP-dsRNA (anti-GFP). We collected ovaries from individual mosquitoes injected with either Norma3-dsRNA or GFP-dsRNA 60h PBM and measured the effect of RNAi on the expression levels of Norma3. Results showed an average expression drop of >50% in anti-Norma3 replicates compared to anti-GFP (Figure 3B), which presented adequate statistical significance (Student’s t-test, P=0.0031) to support our experimental pipeline.

**Figure 3:**
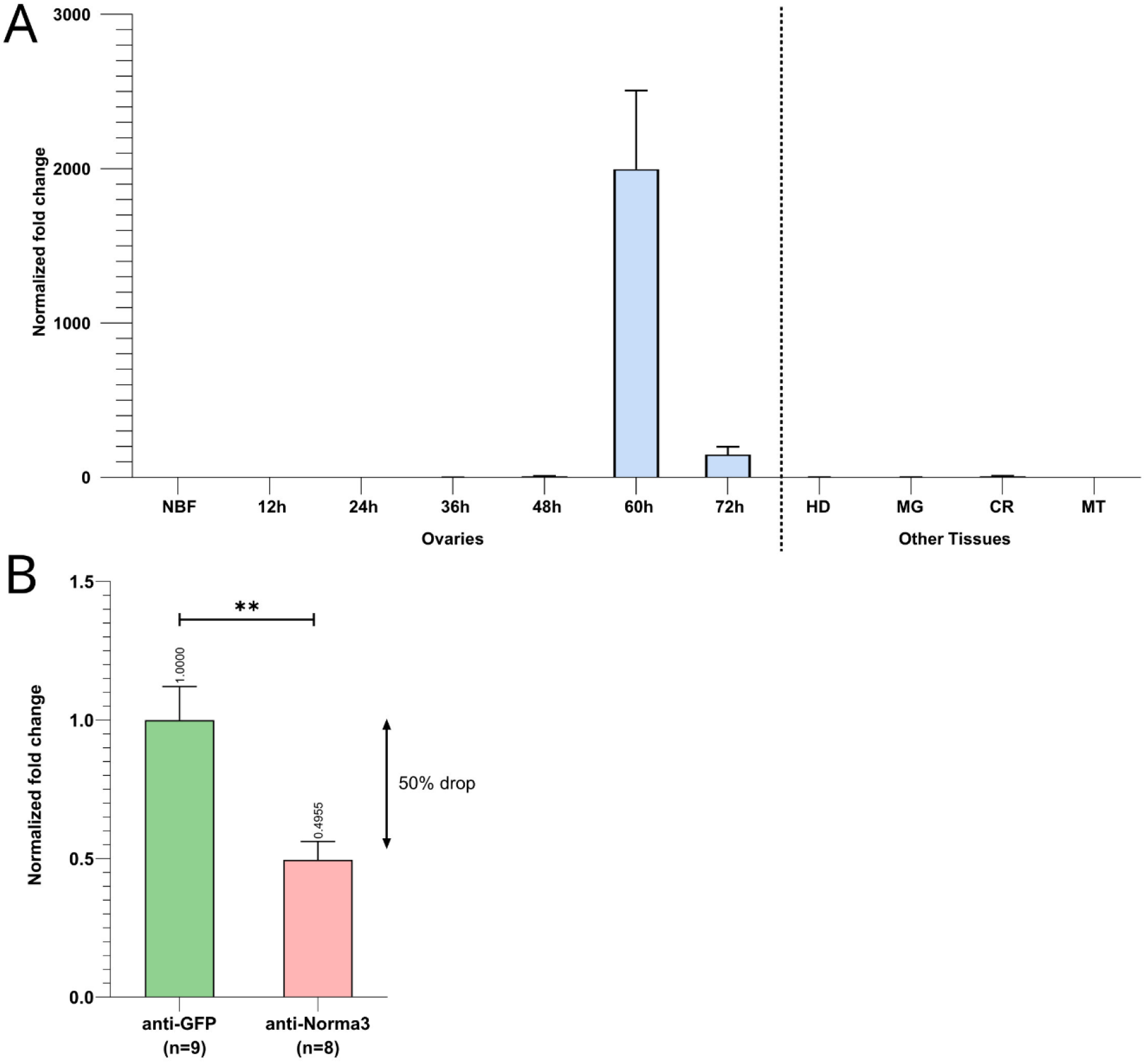
Spatiotemporal expression of Norma3 and its silencing efficacy. **(A)** Expression profile of Norma3 among non-blood fed (NBF) ovaries, post-blood meal (PBM) ovaries and other tissues collected 60h PBM. Ovaries of blood-fed mosquitoes were collected every 12h upon a blood meal, ranging from 12h to 72h PBM when egg development is completed. Ovary samples (NBF & PBM) contained ovaries collected from 5-6 individual mosquitoes (biological replicates). Norma3 exhibits a basal expression in NBF, 12h, 24h, 36h & 48h PBM ovaries, which abruptly increases at 60h PBM leading to a fold change of >2,000 at 60h PBM 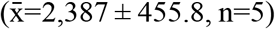 which drops to 200-fold in 72h-PBM 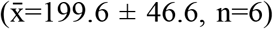, compared to NBF ovaries 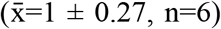. The other tissues that are presented were collected 60h PBM. OV=Ovaries, HD=Head, MG=Midgut, CR=carcass, MT=Malpighian tubes. Each sample contained tissues collected from three individual mosquitoes (biological replicates). Fold change is presented relatively to NBF ovaries. All values were normalized with ribosomal genes RpL32 & RpS17 and are presented as mean ± SEM. **(B)** Relative quantification of Norma3 expression in replicates of the anti-Norma3 dsRNA vs anti-GFP dsRNA samples. Ovaries were collected 60h PBM, the time point when Norma3 peaks its expression. Each sample contains 8-9 biological replicates. Average expression of Norma3 in anti-GFP replicates was set as 1 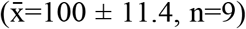 and the overall expression drop of anti-Norma3 replicates, compared to anti-GFP, was measured to 50% 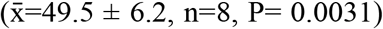. All values were normalized with ribosomal genes RpL32 & RpS17 and are presented as mean ± SEM, **:P-value ≤0.01.

### 3.4 Impact of Norma3 silencing on reproduction

To more deeply characterize the impact of Norma3 silencing on *Ae. albopictus* reproductive ability, we examined various phenotypic traits that are connected to reproduction. First, we looked at the ovary morphology at 60h PBM by measuring the long axis of the ovoid follicle of ovaries dissected by individuals of the anti-Norma3 dsRNA and anti-GFP dsRNA samples. Ovaries of anti-GFP sample presented an average length of 293.6nm, while ovaries of anti-Norma3 presented a significantly lower length of 240nm (Mann-Whitney test, P=0.003) (Figure 4A). Interestingly, anti-Norma3 sample included several much smaller ovaries that had a shorter follicular diameter, while there was evidence of the presence of nurse cells along with their oocyte, indicating their delayed development (Figures 4A-B, Supplementary Figure S1B). Nine anti-Norma3 smaller ovaries that displayed a mean length of 153.4nm were collected and stored for further analysis.

**Figure 4:**
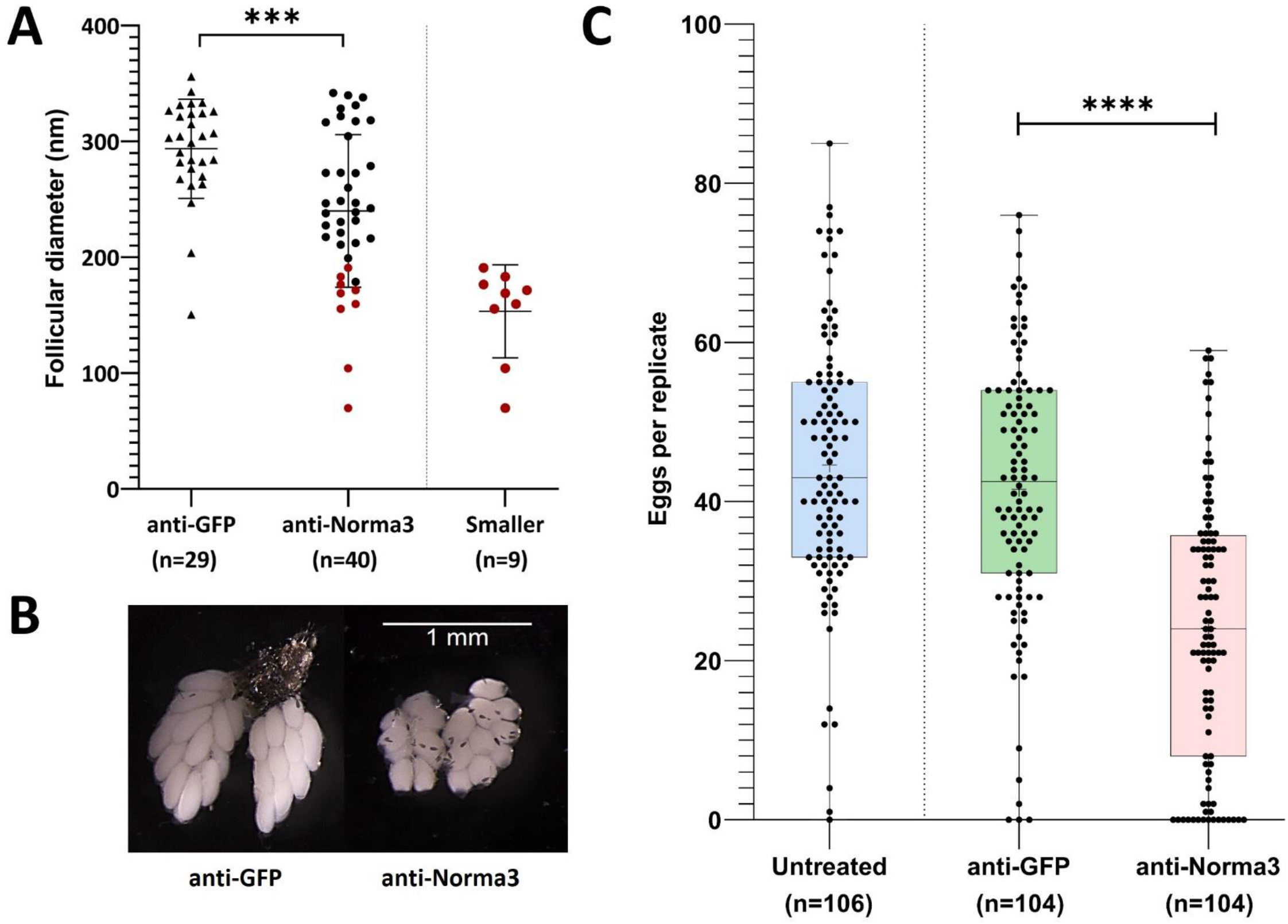
Anti-Norma3 treatment leads to abnormal maturation of ovaries and reduced oviposition. **(A)** Comparison of the length (nm) of the long axis on the ovoid follicles from ovaries obtained from mosquitoes injected with anti-GFP and anti-Norma3 dsRNA. Follicles of the anti-Norma3 sample have a smaller size 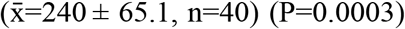, compared to the anti-GFP 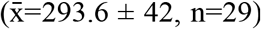. Smaller cohort is a subgroup of the anti-Norma3 sample that includes highlighted smaller replicates 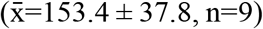. **(B)** Representative ovaries dissected 60h PBM from the anti-GFP and the anti-Norma3 samples. Smaller follicle size and nurse cells are evident in smaller anti-Norma3 ovaries. **(C)** Number of deposited eggs per individual mosquito of untreated 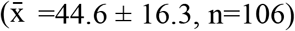, anti-GFP 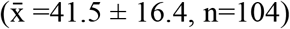 and anti-Norma3 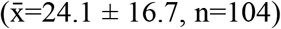 samples. Each dot corresponds to the number of eggs that were laid individually by each female mosquito treated with dsRNA against Norma3 or GFP and untreated control. Each sample contained more than 100 mosquitoes (biological replicates) and an unpaired two-tailed student’s t-test was conducted to assess the statistical significance of the results, by comparing the anti-GFP (control) with anti-Norma3 sample. Anti-Norma3 exhibits statistically significant reduced oviposition. All values are presented as mean ± SD. Error bars include values from min to max. ***: P-value ≤0.001, ****: P-value ≤0.0001

Afterwards, we estimated the effect of the Norma3-dsRNA on oviposition by counting the number of eggs that were laid individually by each female of anti-Norma3, anti-GFP and untreated samples. Mosquitoes of the untreated control laid an average of 44.6 eggs, while mosquitoes of the anti-GFP control laid an average of 41.6 eggs. On the other hand, mosquitoes of the anti-Norma3 treatment laid an average of 24.1 eggs. Interestingly, 14% (n=15) of the anti-Norma3 replicates laid zero eggs, while 26% (n=27) laid less than 10 eggs. The reduction of the average oviposition rate between anti-Norma3 and anti-GFP was 43% and exhibited high statistical significance (Student’s t-test, P<0.0001) (Figure 4C).

Subsequently, we addressed the hatch rate of untreated, anti-GFP and anti-Norma3 samples. We counted the number of larvae that emerged from the eggs that were laid by each individual mosquito of the studied samples and we divided by the total amount of eggs that were laid by each mosquito. Untreated mosquitoes displayed an average hatch rate of 79%, anti-GFP mosquitoes presented a similar hatch rate 78%, while anti-Norma3 mosquitoes had a much lower rate of 37%. (Figure 5A) The difference of the hatch rate between anti-GFP and anti-Norma3 was about 53% and presented high statistical significance (Mann-Whitney test, P<0.0001)

**Figure 5:**
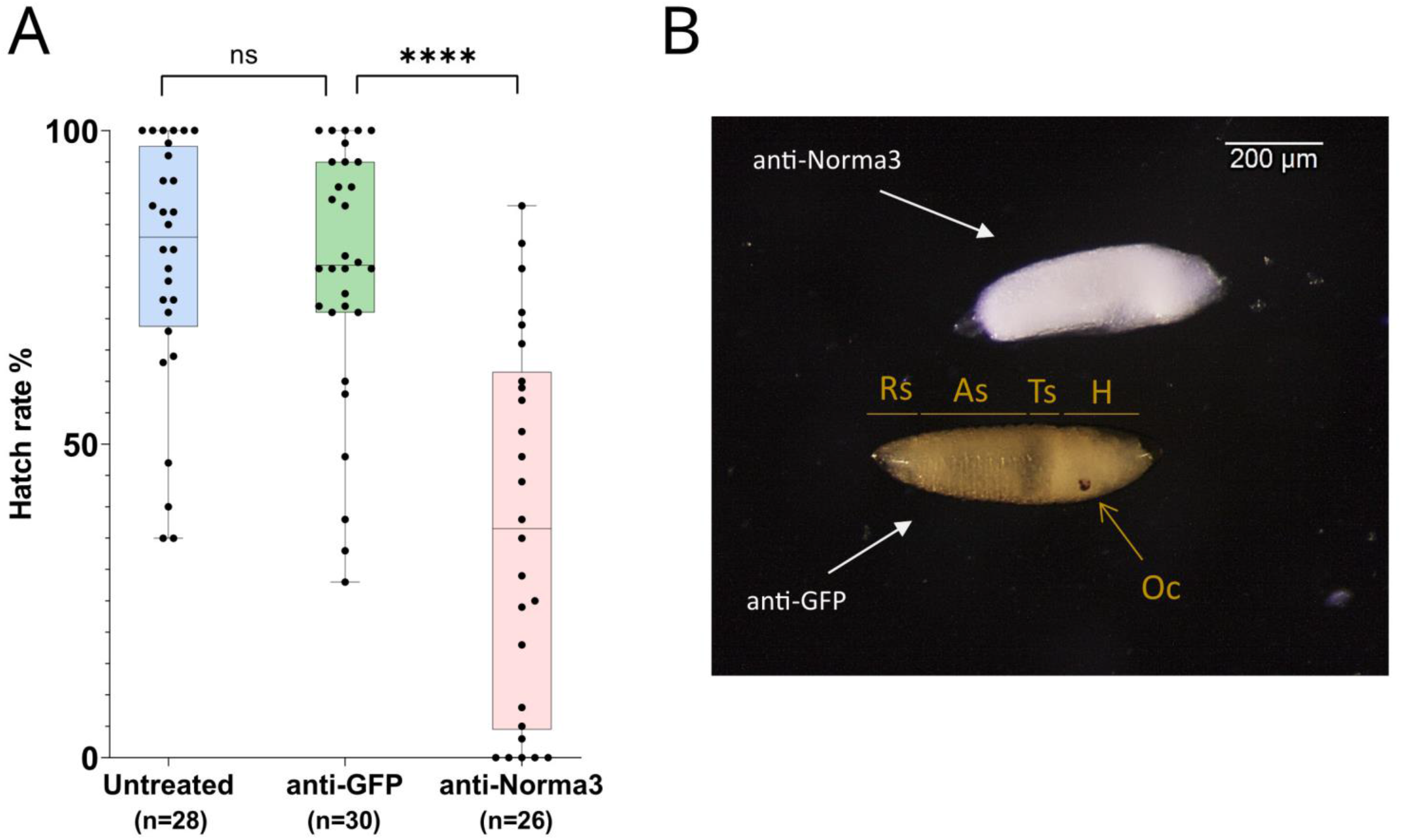
Anti-Norma3 treatment reduces hatch rate and disrupts regular embryonic development. **(A)** Comparison of hatchability of eggs laid by individual females of untreated 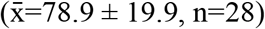, anti-GFP 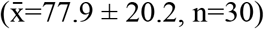, and anti-Norma3 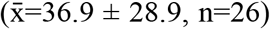 samples. No statistical significance was observed between the untreated and anti-GFP samples, while high significance was detected between anti-GFP and anti-Norma3 samples (P<0.0001, Mann-Whitney test). All values are presented as mean ± SD. ****: P-value ≤0.0001 **(B)** Effect of anti-Norma3 treatment in eggs that were dechorionated with Trpiš solution. Two representative embryos are displayed. Anti-GFP embryo presents regular development as based on the presence of respiratory siphon (Rs), eight abdominal segments (As), thoracic segments (Ts), head (H) and ocelli (Oc). On the contrary, anti-Norma3 does not present any of those structures.

Finally, we attempted to visualize possible larval defects in the anti-Norma3 treated sample. For this, we rendered 72h eggs transparent by bleaching and observed the larvae under the microscope. Significant changes were detected between anti-Norma3 and anti-GFP eggs. In anti-GFP embryos the eight abdominal segments, the thoracic segments, the respiratory siphon, the head and the ocelli were clearly visible indicating regular development of the embryo. On the other hand, anti-Norma3 embryos did not present any of the anticipated normal structures; instead, they exhibited a defective appearance of an undifferentiated mass (Figure 5B, Supplementary Figure S2B).

### 3.5 Regulatory interplay of Norma3 with neighboring mucins

Since lncRNAs often regulate coding genes in their genomic neighborhood (Rinn and Chang, 2012; Engreitz et al., 2016; Joung et al., 2017; Khyzha et al., 2019; Xing et al., 2021), we investigated the possible association of Norma3 with protein-coding genes in its vicinity. We returned to the available transcriptomic study containing NBF and PBM ovary datasets (Gamez et al., 2020) and assessed the correlation of Norma3 expression against the rest of the transcriptome. Within the region from 100kb upstream to 100kb downstream of Norma3, we identified 8 annotated transcripts (i.e., 4 mucins, 1 venom protein and 3 chymotrypsin inhibitors) which all exhibited robust positive correlation with Norma3 (Pearson correlation coefficient > 0.96, maximum FDR = 2.17e-9) (Supplementary Data S2). Subsequently, we reduced the transcriptome to transcripts which were more likely to present any change in ovaries among early (i.e., 12h-24h), middle (36h-48h) and late (60h-72h) time points (posterior-probability EBSeq > 75%). This transcript set (n = 10314) was subjected to clustering analysis based on their expression over the entire time course (i.e., NBF, 12h, 24h, 36h, 48h, 60h and 72h PBM). Norma3 was grouped together with 114 other genes (111 protein-coding and 3 long non-coding) in a cluster of genes that was not expressed in NBF and early PBM samples but initiated low transcription at 36h and peaked at the end of vitellogenesis (around 48h PBM) (Figure 6A). Among the 8 positively correlated neighbor transcripts, three mucins belonged to a cluster that presented an intense expression peak at 48h (Figure 6B).

**Figure 6:**
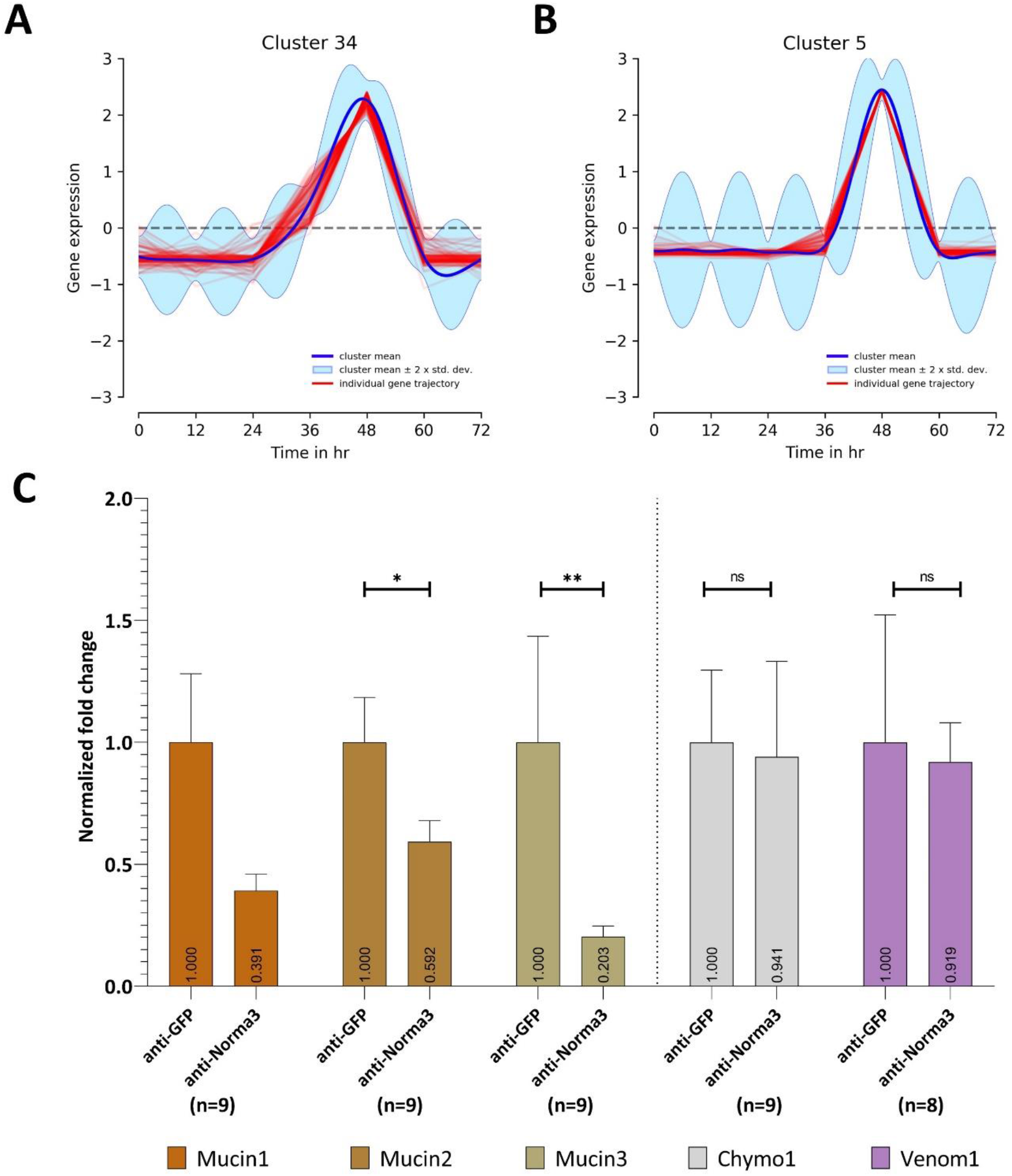
Expression patterns through post blood meal timepoints (A-B) and in relation to Norma3 silencing (C). Based on its expression across all timepoints, Norma3 was grouped in Cluster 34, which gradually peaks at 48h post blood meal. **(B)** Three mucins neighboring Norma3 (mucin1-3) belong to Cluster 5 which presents an acute peak at 48h. **(C)** Expression of Norma3-neighboring proteins in replicates of the anti-Norma3 vs anti-GFP sample. All mucins exhibit a statistically significant expression drop. Mucin1 61% 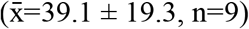, mucin2 41% 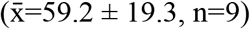, mucin3 80% 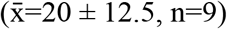 while other neighboring proteins (chymo1, venom1) do not (Mann-Whitney test). Both anti-GFP and anti-Norma3 samples contain 9 biological replicates. Results were normalized with ribosomal genes RpL32 & RpS17. All values are presented as mean ± SEM. *: P-value ≤0.05, **: P-value ≤0.01.

We then generated their detailed expression profile in NBF and PBM ovaries. We confirmed the expression pattern of five of the neighboring transcripts via qPCR. Three were annotated as mucin-2 and mucin-2-like (mucin1, 2, 3), one was annotated as cysteine-rich venom protein 6-like (venom1) and one as chymotrypsin inhibitor (chymo1). All five genes presented basal expression in NBF, 12h, 24h, 36h & 48h ovaries. The expression of the three mucins presented a sharp increase between 12,000 (mucin2) and ∼40,000-fold (mucin3) in 60h PBM ovaries, while chymo1 increased ∼10,000-fold in 72h PBM and venom1 3,000-fold in 60h PBM ovaries (Supplementary Figure S3). To further assess the potential impact of Norma3 on the mucins we assayed the expression of the three mucins in a small sample of smaller ovaries (Figure 4A) of the anti-Norma3 60h-PBM sample (that we collected as described in section 3.4.). Statistically significant downregulation between 40% to 80% of the three mucin genes was detected in the smaller anti-Norma3 ovaries compared to anti-GFP control (Figure 6C). Downregulation of mucin3 presented the most significant effect, while no significant effect was detected on other protein-coding transcripts located in the same genomic region (chymo1, venom1). Nevertheless, since we only tested nine smaller anti-Norma3 ovaries, we acknowledge that this result serves as a preliminary indication of the potential impact of Norma3 on mucin expression that should be validated with larger datasets.

## 4 DISCUSSION

Long non-coding RNAs have arisen during the last decades as a fascinating field of research due to their unique features and intriguing mechanisms of action in the absence of a protein product. One entirely unexplored field of potential lncRNA applications is pest management and mosquito control. Due to the very low sequence conservation among lncRNAs (Diederichs, 2014; Tavares et al., 2019), such genes could be ideal as species-specific targets in new generation genetic control approaches. However, up to date there has not been sufficient research on specific lncRNAs to serve as efficient molecular targets with potential utility in pest management. In the tiger mosquito *Ae. albopictus*, a competent vector of multiple arboviruses, only a few studies regarding ncRNAs have been published so far (Gu et al., 2019; Azlan et al., 2021; Betting et al., 2021) and none presented significant data related to the impact of particular lncRNAs in physiological systems. Our aim was to provide the first proof-of-evidence regarding the role of specific lncRNAs in a vital biological process of the mosquito with potential utility for pest control. We focused on the reproductive system because of its significance for mosquito’s viability and the broad applications that it may offer in mosquito control.

After computationally annotating predicted lncRNAs regarding their coding potential, genomic localization, species and developmental context specificity (Figure 1), we identified 10 species-specific lncRNAs which were overexpressed in *Ae. albopictus* ovaries upon blood-feeding (Figure 2A) and set out to further explore their potential role in reproduction. We deployed a loss-of-function RNAi-mediated pipeline and investigated the changes that occurred in the fecundity of female mosquitoes upon silencing of each lncRNA (Figure 2B). We focused on an antisense intergenic lncRNA, that we named Norma3, because its targeting provoked a robust phenotypic effect. Moreover, it exhibited an ideal expression profile: basal expression in most of the NBF/early PBM time points followed by a sharp increase by the end of vitellogenesis (Figure 3A). Its expression was also highly ovary-specific (Figure 3A), while its nucleotide sequence did not display any similarities to annotated genes of other species. It is worth mentioning that we observed a slight discordance between expression peaks of RNA seq data and qRT-PCR, probably due to different sampling methodologies or adaptation of the local *Ae. albopictus* strain to our laboratory conditions.

Our successful gene silencing approach (Figure 3B) resulted in significant reduction in oviposition and egg hatching (Figures 4C, 5A) of the anti-Norma3 dsRNA-treated mosquitoes. We further associated this fertility reduction with defective ovaries (Figure 4B), smaller ovary follicle size (Figure 4A) and obvious malformations of 72-hour embryos of the anti-Norma3 treated mosquitoes (Figure 5B). In addition, we attempted to obtain insights on the potential mode of action of Norma3. Long non-coding RNAs operate in a variety of different modes and uncovering their functional role is not a trivial task. Unlike protein-coding genes that share conserved domains due to sequence homology, the features of lncRNAs mainly arise from their secondary structures which perform complex interplay with DNA regions, RNAs or proteins (Pang et al., 2006; Ding et al., 2014). One of most frequent roles of lncRNAs is to regulate the expression of coding genes and, oftentimes, such genes are localized in the vicinity of a lncRNA (Rinn and Chang, 2012; Engreitz et al., 2016; Joung et al., 2017; Khyzha et al., 2019; Xing et al., 2021). To explore this possibility, we performed *in silico* analysis to highlight coding genes that possessed similar expression profiles as Norma3 and were localized in the same region ±100kb up-or-down stream of Norma3 (Supplementary Data S2). Our search resulted in eight coding genes which were annotated as mucins, chymotrypsin inhibitors and cysteine-rich venom-like proteins, while all of them contained a trypsin inhibitor-like (TIL) cysteine rich domain. According to our qPCR results, the expression of three of these mucins was severely diminished by 70% upon injection of anti-Norma3 dsRNA (Figure 6C). While this evidence is circumstantial, it may indicate a possible interplay between Norma3 and the neighboring mucins. Mucins are proteins that are characterized by domains that contain repetitive sequences of proline, threonine and serine (PTS domains) and heavy O-glycosylation (Tran and Ten Hagen, 2013) which are present in most metazoan (Lang et al., 2007). While mucins are the principal components of mucus and mucous membranes, they carry diverse roles from lubrication to cell signaling to forming chemical barriers.

To our knowledge, there are no direct studies associating mucins with oviposition in mosquitoes. Ovary-specific mucins have been highlighted in *Ae. albopictus* by a recent study that reported the upregulation of multiple mucins upon blood-feeding in females and speculated a potential role of these genes in oviposition (Deng et al., 2020). Nonetheless, various ovary-specific mucins have been identified in other insects and studies have revealed their role in ovary development and reproduction. In particular, mucin-like proteins have been associated with the eggshell, a multilayered structure which is formed during oogenesis that protects and nurtures the developing embryo prior to its arrest (Osterfield et al., 2017). In *Drosophila melanogaster* three putative eggshell genes code for proteins with mucin-like domains (Muc4B, Muc11D, Muc12Ea). Muc4B has been suggested to be a component of the wax layer of the embryo, while Muc11D and Muc12Ea potentially act as mediators of chorion hardening and coating for passage of the embryo through the oviduct (Tootle et al., 2011). Another ovary-specific mucin (NlESMuc) that was identified in plant grasshopper *Nilaparvata lugens* was also related to the eggshell and was proven essential for its fecundity. Specifically, the RNAi-mediated targeting of NlESMuc caused reduced oviposition, lower egg production and less egg hatching (Lou et al., 2019). Finally, a study in the lepidopteran *Spodoptera exigua* presented an ovary-specific mucin-like protein called Se-Mucin1 that was associated with choriogenesis. In the absence of Se-Mucin1, females exhibited reduced fecundity and the hatch rate of the eggs was also significantly impaired, while SEM analysis of the eggshell structures revealed that they were remarkably malformed (Ahmed et al., 2021).

Through our analysis we collected pieces of circumstantial evidence that indicates a possible interplay between Norma3 and three neighboring mucins (mucin1, 2, 3). Firstly, both RNA seq analysis (Fig. 6A-B) and qPCR data (Fig. 3A, Supplementary Fig. S3) demonstrate tightly linked sharp expression increases of Norma3 and the three mucins at 60 hours post blood meal. Moreover, we detected a potential influence of anti-Norma3 treatment in the expression drop of mucins 1, 2 & 3 that could be associated to the developmental delay of ovaries (Fig. 6C). Anti-Norma3 treatment led to a downregulation of their expression, especially in mucin3 which exhibited the most intense and statistically significant effect (P-value: 0.0019). However, the current sample size of anti-Norma3 and anti-GFP ovaries should be enlarged in order to validate the finding. Given these preliminary results, we speculate that Norma3 may act as a positive regulator of the mucin cluster and its silencing could lead to inhibition of their ovary specific-expression and possibly to disruption of the reproductive ability of *Ae. albopictus. Cis*-acting lncRNAs are one of the most dominant lncRNA classes and the majority of them overlap with enhancers elements (Gil and Ulitsky, 2020). We also presume that Norma3 and the mucin proteins are involved in the formation of the eggshell, a hypothesis which is based on the well-studied role of mucins in other insects (Tootle et al., 2011; Osterfield et al., 2017; Lou et al., 2019; Ahmed et al., 2021), but also relies on the relevant phenotypic outcomes that were provoked by targeting an eggshell-related protein (EOF-1) in the relative species *Ae. aegypti* (Isoe et al., 2019). Further research is necessary to verify the hypotheses on both the impact of mucins on mosquito reproduction and their regulatory association with Norma3.

Our study presents a conceptual pipeline and a proof-of-principle towards novel approaches of insect pest control. It begins with the discovery of lncRNAs involved in the regulation of a physiological system that is fundamental for species survival and propagation, the reproductive system, and it showcases that down-regulating a particular lncRNA results in the damage of that system, reducing insect fecundity and fertility. Our results suggest that species-specificity of lncRNAs renders them preferred targets of RNAi-based pesticides. In fact, RNAi technology has offered a new and hopeful prospect of ecologically friendly approaches to insect control since it can minimize off-target effects. Low sequence conservation and high species-specificity of lncRNAs provide extra added value towards that end. Nonetheless, delivery methods of RNAi-based insecticides still pose major challenges (Yu et al., 2013; Niu et al., 2018) Till now, transgenic plants producing dsRNAs against vital insect genes has been the most straightforward and efficient application of RNAi technology for insect control (Nowara et al., 2010). However, this approach entails public acceptance problems due to the transgenic nature of the plants (Herman et al., 2021). Alternatively, spraying of stabilized dsRNA is recently being considered and actively researched [reviewed in (Rank and Koch, 2021)]. Examples come for the use of this technology against plant pests [e.g., sprays of dsRNA-producing *E. coli* against 28-spotted ladybird (Wu et al., 2021) or dsRNA sprays against Colorado potato beetle (Mehlhorn et al., 2020)], but one can easier envision house sprayings against biting mosquitoes due to greater dsRNA stability inside a house environment. Furthermore, reproductive system related lncRNAs, such as Norma3, could also be exploited in cutting-edge gene drive approaches aiming to suppress disease vectors by reducing female fertility (Hammond et al., 2016; Kyrou et al., 2018; North et al., 2020; Simoni et al., 2020). Given the available sequencing data in several insect species of public health or agricultural importance (or the affordability of obtaining such data from any organism of choice), this pipeline can be adopted to any given species and yield novel species-specific targets for pest control, thus addressing one of the most difficult challenges of the pesticide industry for species-specificity and environmental safety.

## Supporting information

Supplementary Data S1. LncRNA annotation

Supplementary Data S2. Norma3 correlation

Supplementary Data S3. Experimental values

Supplemental Table S1. Gene names

Supplemental Table S2. Primers

## 5 ACKNOWLEDGMENTS

We thank Prof. N. Papadopoulos and Dr. C. Ioannou of the Dept of Agriculture, Crop Production and Rural Environment, University of Thessaly, for kindly providing the mosquito strain used in the present study and Assist. Prof. S. Vasileiadis from the Dept. of Biochemistry & Biotechnology, for his guidance on statistical analyses.

