## Supplementary figures for "A species-specific lncRNA modulates the reproductive ability of the Asian tiger mosquito"

### *Supplementary Material*

**A)**

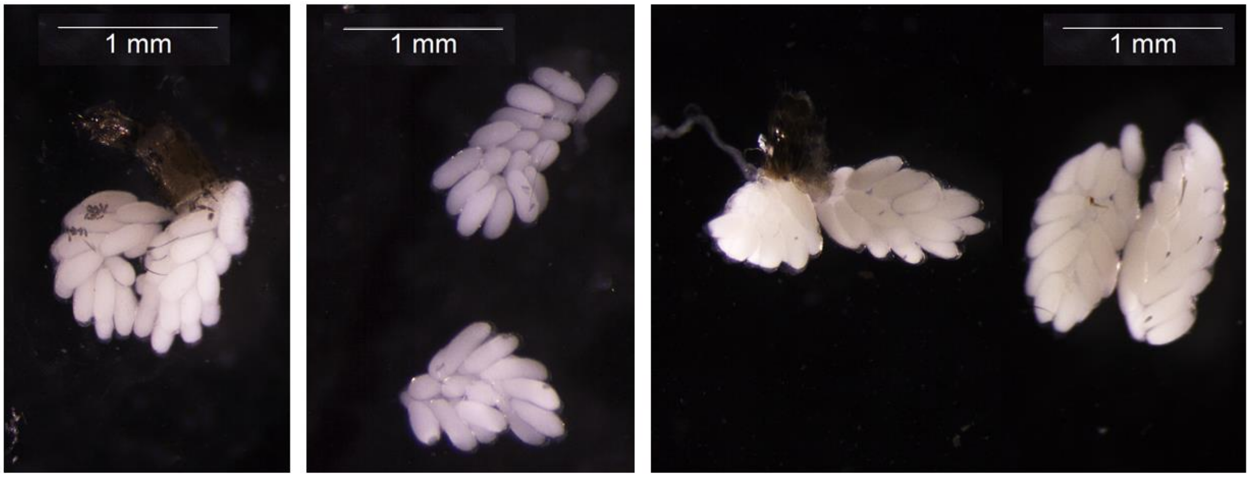


**B)**

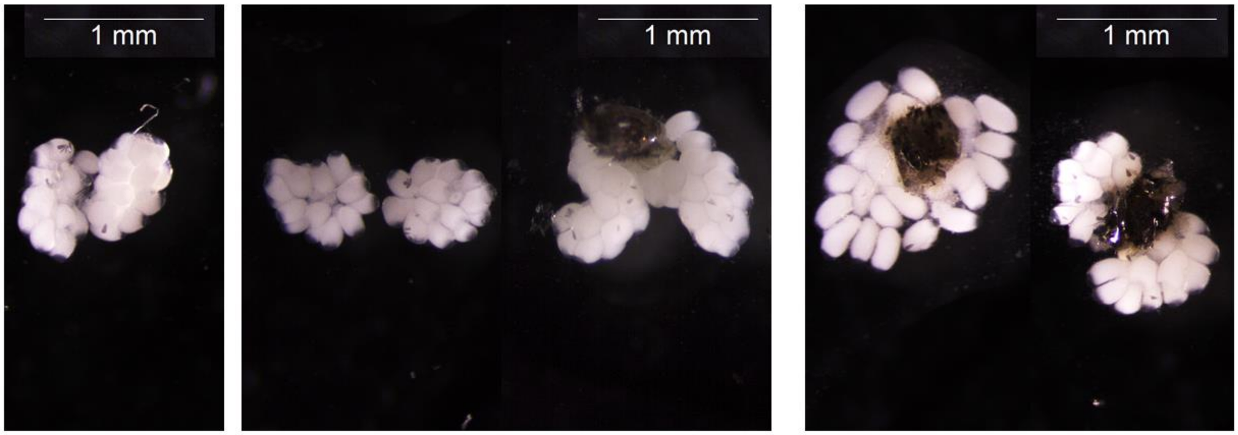


**Supplementary Figure S1:** Microscope images that represent ovaries dissected 60h post-blood meal from mosquitoes that belonged to (A) anti-GFP dsRNA sample, (B) anti-Norma3 dsRNA sample.

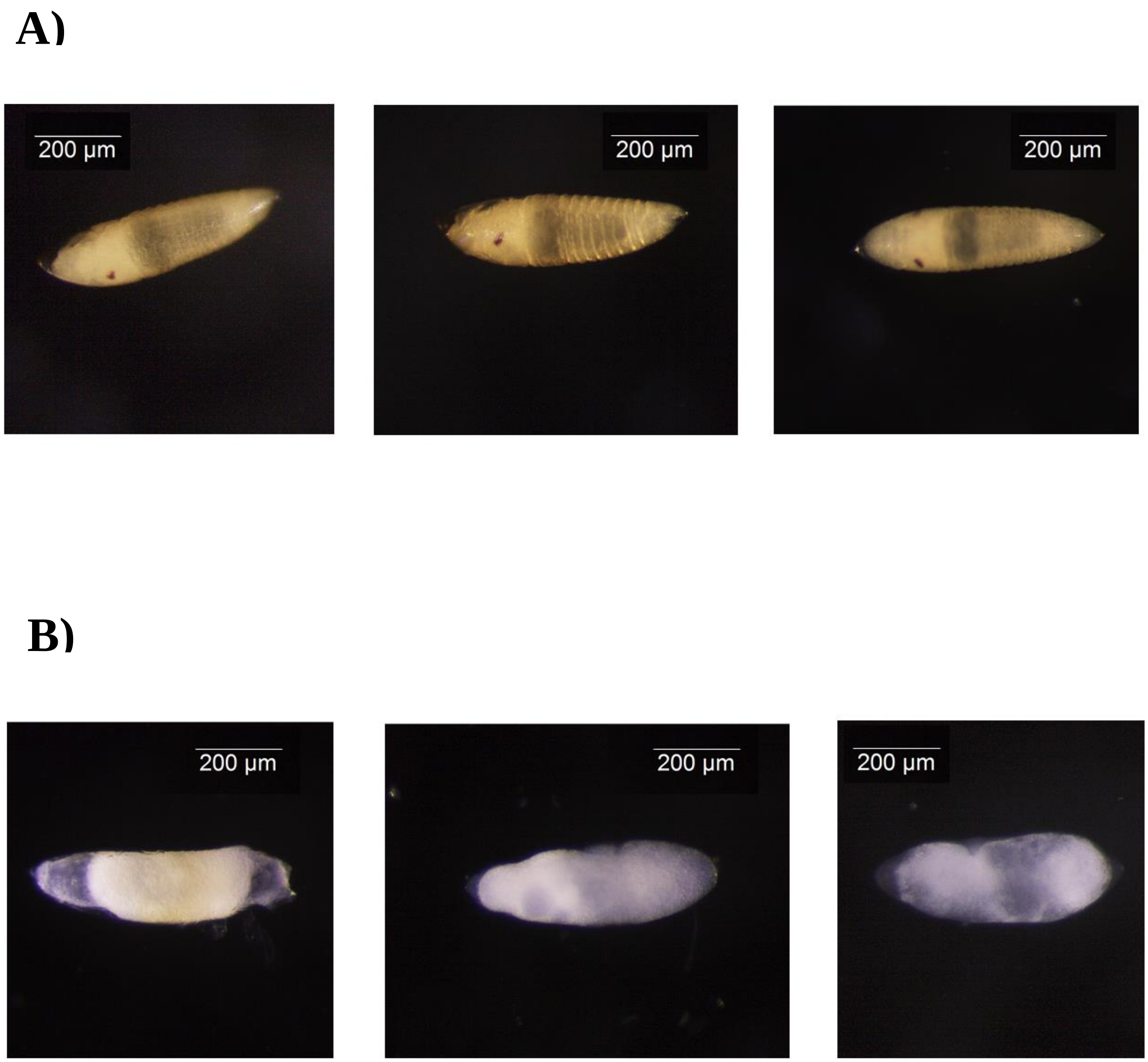


**Supplementary Figure S2:** Microscope images that represent embryos that were dechorionated 72h post egg-laying from mosquitoes that belonged to (A) anti-GFP dsRNA sample, (B) anti-Norma3 dsRNA sample.

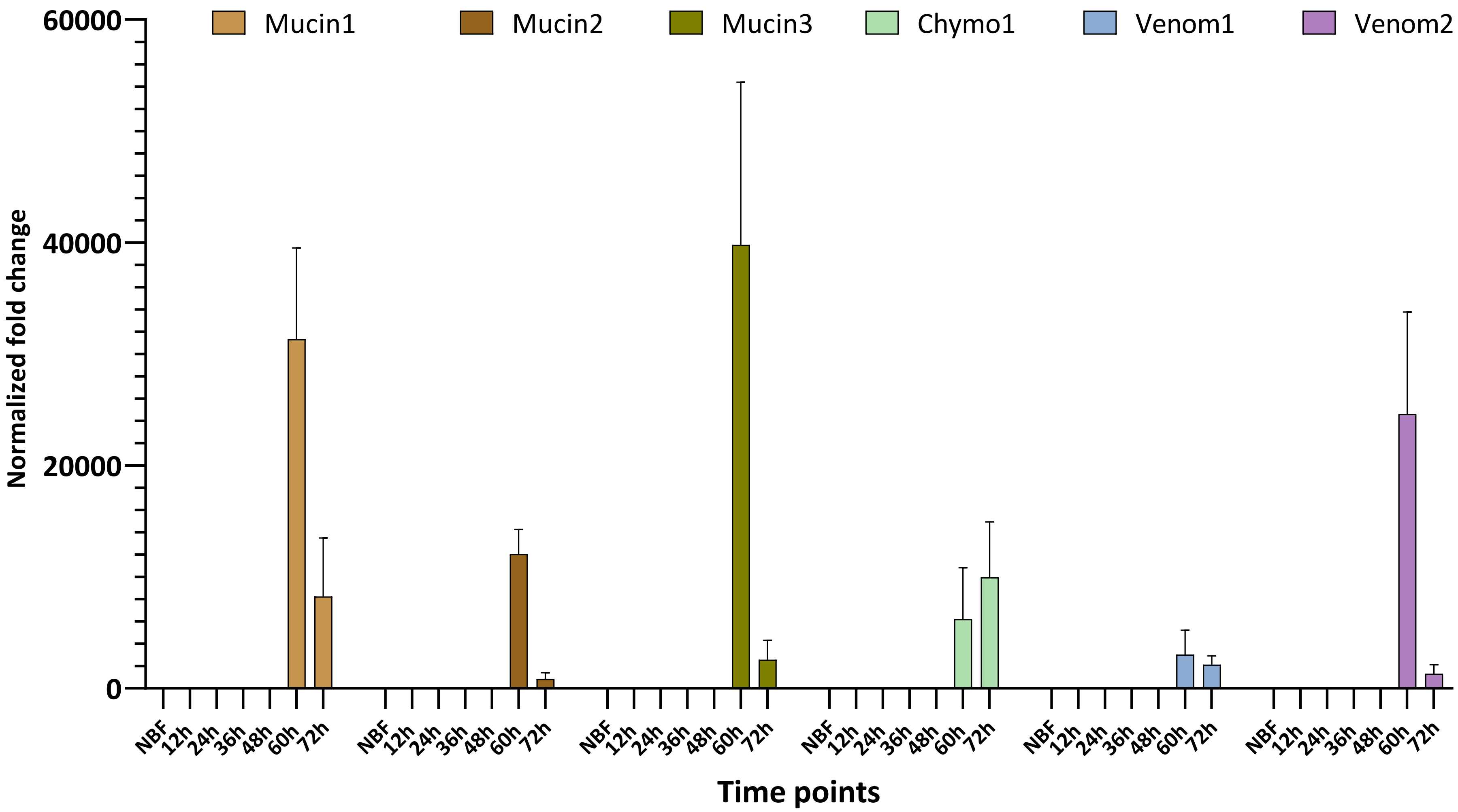


**Supplementary Figure S3**. Expression profile of 5 protein-coding genes in ovaries of non-blood fed (NBF) and post blood-meal (PBM) mosquitoes in serial time points. The presented genes are positively correlated to Norma3 and are located in genomic vicinity. All expression values correspond to normalized fold change of each sample relatively to NBF and are presented as mean ± SEM. Results were normalized with ribosomal genes RpL32 & RpS17. Each group contained 3 biological replicates, except of 60h & 72h groups which included 6 biological replicates.
